# Mixed Models for Meta-Analysis and Sequencing

**DOI:** 10.1101/020115

**Authors:** Brendan Bulik-Sullivan

## Abstract

Mixed models are an effective statistical method for increasing power and avoiding confounding in genetic association studies. Existing mixed model methods have been designed for “pooled” studies where all individual-level genotype and phenotype data are simultaneously visible to a single analyst. Many studies follow a “meta-analysis” design, wherein a large number of independent cohorts share only summary statistics with a central meta-analysis group, and no one person can view individual-level data for more than a small fraction of the total sample. When using linear regression for GWAS, there is no difference in power between pooled studies and meta-analyses [1]; however, we show that when using mixed models, standard meta-analysis is much less powerful than mixed model association on a pooled study of equal size. We describe a method that allows meta-analyses to capture almost all of the power available to mixed model association on a pooled study without sharing individual-level genotype data. The added computational cost and analytical complexity of this method is minimal, but the increase in power can be large: based on the predictive performance of polygenic scoring reported in [2] and [3], we estimate that the next height and BMI studies could see increases in effective sample size of ≈15% and ≈8%, respectively. Last, we describe how a related technique can be used to increase power in sequencing, targeted sequencing and exome array studies.

Note that these techniques are presently only applicable to randomly ascertained studies and will sometimes result in loss of power in ascertained case/control studies. We are developing similar methods for case/control studies, but this is more complicated.

## Notation

We use the following notation throughout:

- *N ∈* ℕ: sample size.
- *y ∈* ℝ^*N*^: centered and standardized phenotype.
- *X*_*j*_ *∈* ℝ^*N*^: centered and standardized vector of genotypes at the test variant *j*.
- *β*_*j*_ *∈* ℝ: effect size of the test variant *j*.
- ***X***_*−j*_: *M × N* matrix of centered standardized genotypes excluding a region around *j*.

As in [4], we assume that all fixed effects (*e.g.,* age, sex, batch, study indicators, principal components of the genotype matrix *etc*) have already been projected out of the data. The meta-analysis method proposed here achieves protection from population stratification and proper calibration with related samples using BOLT-LMM. In order to simplify notation, we omit most discussion of population stratification and family structure; these issues have already been addressed with regard to BOLT-LMM in ref [4].

Our goal is to test the null hypothesis ℋ_0_: *β*_*j*_ = 0.

## Association Testing with Linear Regression

The simplest approach to testing genetic variants for association with a quantitative phenotype is to fit the model

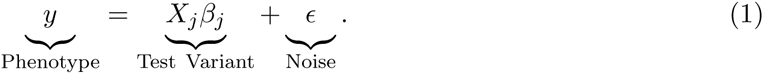

using linear regression (typically we would also include the top several principal components of the genotype matrix as covariates; as previously noted, we assume without loss of generality that these have already been projected out). Linear regression yields an effect-size estimate *β*_*j,*lin_ and a variance estimate 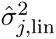 We can construct χ^2^ statistics

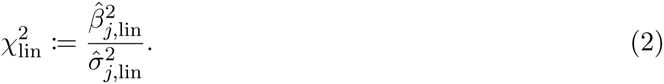

This test statistic is called the Armitage Trend Test (ATT) statistic. The choice of linear regression statistic is not particularly important; all of the standard linear regression test statistics are asymptotically equivalent. The ATT statistic asymptotically follows a one degree-of-freedom noncentral χ^2^ distribution. For small *β*_*j*_ (typical in GWAS), 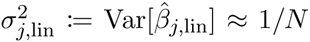, so the noncentrality parameter is

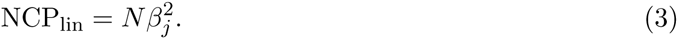

Note that since *β*_*j*_ is the standardized effect size of variant *j*, 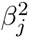 is variance explained.

## Association Testing with Mixed Models

The linear regression model from Equation 1 treats the effects of all variants other than the test variant as noise. We can increase the power of the test for association by explicitly modeling the effects of the other variants. The model then becomes

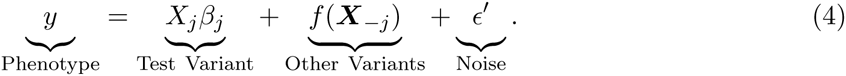

Note that the noise term *∊*^*′*^ in Equation 4 has smaller variance than the noise term *E* in Equation 1, because *∊* = *∊*^*′*^ + *f* (***X***_*-j*_). If we allow *f* to take arbitrary functional form, then this is *additive* mixed model; a linear mixed model is the special case where we constrain *f* to be a linear function. This model is fit using a two-step procedure [4]:

1. Generate predictions 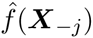 and prediction residuals 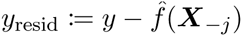.
2. Obtain an effect-size estimate and variance 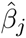 and *s*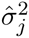 by fitting the model *y*_resid_ = *X*_*j*_*β*_*j*_ + *E* using linear regression.

Refs [4, 5] refer to step 1 as ‘de-noising’ the phenotype. It is important to use predictions generated with the test variant and variants in LD with the test variant left out. By including the effect of the test variant in the predictions, we remove most of the effect of the test variant from the residual phenotype, which results in loss of power in step 2. Ref [6] coined the term ‘proximal contamination’ to describe this problem. We can avoid proximal contamination by generating predictions with the test variant and its LD partners removed (for example, ref [4] train 22 predictors, leaving out one chromosome at a time).

The mixed model fitting algorithm can be used with arbitrary choice of predictor 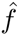. A large number of different mixed model association methods have been published in the statistical genetics literature, and the distinctions between most of these methods can be understood as different statistical and algorithmic choices for fitting 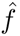 (at least in the special case where individuals in the study are unrelated). For example, GCTA generates predictions using BLUP [5], which is equivalent to ridge regression with the shrinkage parameter set via maximum-likelihood. FAST-LMM [7] uses faster algorithms to fit the same model (GEMMA [8] and EMMAX [9] also fit the same model, but without avoiding proximal contamination and so suffer from loss of power [5]). FAST-LMM-Select [6] uses ridge regression with a feature selection step. LMM-LASSO [10] uses a Lasso with shrinkage parameter selected via cross-validation. BOLT-LMM [4] uses Bayesian linear regression with a mixture-of-Gaussians prior with hyperparameters selected via a combination of (stochastic) maximum-likelihood and cross-validation. This list of examples is not intended to be exhaustive, but illustrates the general principle.

The variance of the standardized effect size estimate 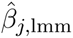 from the mixed model applied to a single (pooled) study is 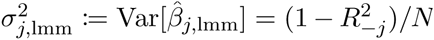, where 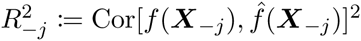 is the prediction *R*^2^ achieved by the mixed model predictor (leaving out a region around variant *j* in order to avoid proximal contamination). Thus, we can construct mixed-model *χ*^2^ statistics

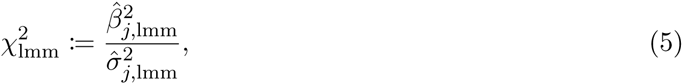

which asymptotically follow a noncentral one degree-of-freedom *χ*^2^ distribution with noncentrality parameter

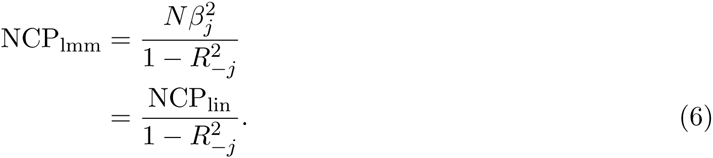

Note that the power increase depends on the *true* prediction *R*^2^, *i.e.,* the squared correlation between 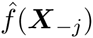 and *f* (***X***_*-j*_) (which is not observable) [4]. The training sample prediction *R*^2^, *i.e.*, 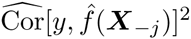, will typically over-estimate the true prediction *R*^2^, because error metrics on the training set are optimistic. For power calculations, one can obtain a better estimate of 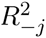 using cross-validation, as in [4].

The *Χ*^2^ statistic from Equation 5 is equivalent to the retrospective quasilikelihood score statistic from BOLT-LMM in the case where Var[*X*_*j*_] = ***I***, *i.e.,* for GWAS that sample unrelated individuals (ref [4], Supplementary Equation 22). Association testing in GWAS datasets with family relatedness is more difficult; in particular, residualizing on predictions generated from typed SNPs (equivalently, using the genetic relatedness matrix estimated from typed SNPs) achieves suboptimal power and suffers from inflated type I error due to residual non-independence of family members (*e.g.,* from correlated environmental effects, or the effects of untyped rare variants). For an overview of mixed model methods applied to family data, see ref [11].

## Meta-Analysis Terminology and Study Design

For readers unfamiliar with the analysis procedure used by consortia such as GIANT, we provide a brief review of GWAS meta-analysis.

Gathering large samples of genotype-phenotype data is difficult, time-consuming and expensive. As a result, most GWAS datasets are generated not by a single lab, but rather by large consortia that pool the efforts of many research groups working in parallel. We refer to the data generated by an individual research group as a ‘cohort’.

One approach to GWAS is to aggregate the genotype and phenotype data from all cohorts onto a single computer, then to run regressions on the combined dataset. We refer to this as a ‘pooled’ study design. Some examples of consortia that use the pooled design include the Psychiatric Genomics Consortium [12–16] and the International Inflammatory Bowel Disease Consortium [17].

Studies that use the pooled design represent a minority of all GWAS, because there are often restrictions (imposed by national law, IRB regulations, etc) that prohibit researchers from sharing individual-level genotype data with other groups. When researchers cannot share individual-level genotype data, an alternative approach is for each research group to run regressions within their own cohort, then for the research groups to share summary statistics (effect size estimates and variances, or equivalent) with a central meta-analysis group, who then meta-analyze the summary statistics using the methods described in the next section. We refer to this study design as a ‘meta-analysis’.

## Inverse-Variance Meta-Analysis

Suppose we have summary-level association results from *S* non-overlapping studies of the same phenotype, where the summary data consist of a (standardized) effect-size estimate 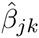 and a variance estimate *s*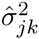 for *k* = 1*, …, S*. We suppose that the 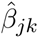 are consistent estimates of a single underlying parameter *β*_*j*_, and that the *s*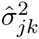 are consistent estimates of the true sampling variances 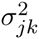. For asymptotically normal estimators (a class which includes all both the linear regression and mixed model estimators described here), the sampling distribution is 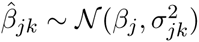. In GWAS, the standard approach for combining these data is inverse-variance meta-analysis [18], which is the minimum-variance unbiased estimator of the true effect *β*_*j*_ The inverse variance meta-analysis effect size and variance estimates are

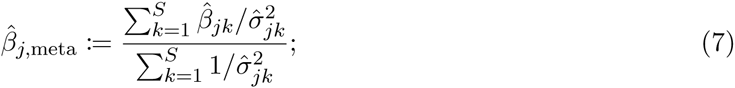

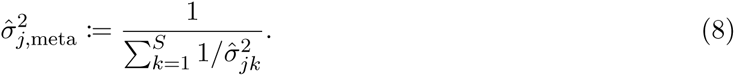

The asymptotic sampling distribution is therefore *β*_*j*,meta_ 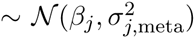. The meta-analysis *Χ*^2^ statistic is

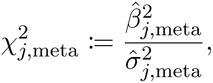

which follows a one degree-of-freedom noncentral *Χ*^2^ distribution with noncentrality parameter

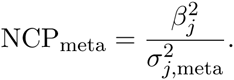

Here, 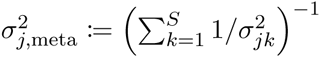 is the true (rather than estimated) meta-analysis variance.

## Meta-Analysis Noncentrality Parameters

The noncentrality parameter of an inverse-variance weighted meta-analysis of linear regression association tests is

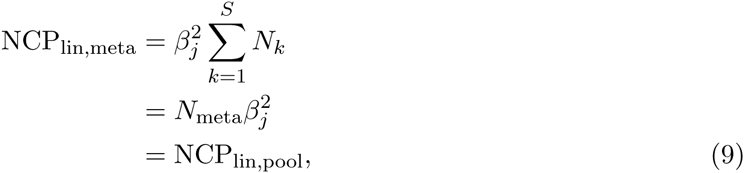

where *N*_meta_:= *k N*_*k*_, and NCP_lin,pool_ denotes the noncentrality parameter of linear regression applied to a pooled study with sample size *N*_meta_. Hence, when using linear regression for association testing, there is no difference in power between a pooled study and a meta-analysis of equal size [**?**].

The noncentrality parameter of an inverse-variance weighted meta-analysis of mixed model association tests is

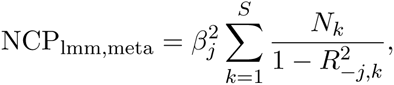

Where 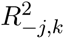 denotes the prediction *R*^2^ achieved by the mixed model predictor leaving a region around variant *j* out trained using only the samples from study *k*. The previous expression is exact, and should be preferred for power calculations. In order to obtain a simpler expression for intuition, we make the approximation 1*/*(1 − *R*^2^) ≈ *R*^2^ + 1 (which holds for small *R*^2^). This allows us to (approximately) simplify the previous expression to

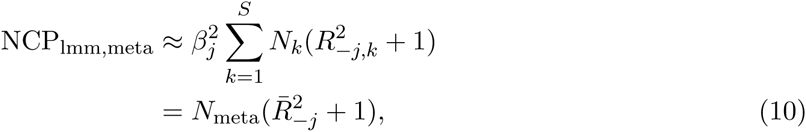

where

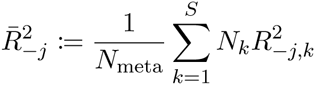

is the sample-size weighted average prediction *R*^2^ achieved by mixed model predictors trained using data from one cohort at a time (leaving out a region around the test variant *j* in order to avoid proximal contamination). In contrast, the noncentrality parameter of mixed model association applied to a pooled study of equal size is

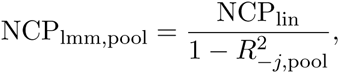

where 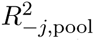 denotes the prediction *R*^2^ achieved by the mixed model predictor trained on the pooled sample (again, leaving out the region around variant *j*). In a typical application, *N*_meta_ might be on the order of hundreds of thousands, while *N*_*i*_ will be one to two orders of magnitude smaller. Since a predictor trained on hundreds of thousands of individuals will perform much better than a predictor trained on tens of thousands of individuals, 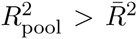. Thus, the relationship between the four noncentrality parameters described above is

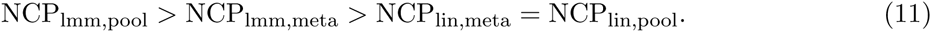

## More Powerful Strategies for Mixed Model Meta-Analysis

The origin of the loss of power in mixed model meta-analysis compared to pooled mixed model association is the poor performance of predictors trained only on individual cohorts. Therefore, what is required is a method for training a predictor on the whole dataset without sharing genotype data; that is, a method for training a predictor using the summary statistics. Conveniently, such methods have already been described [19–21]. At the time of writing, the most sophisticated general-purpose method in this class appears to be LDpred [19]; though we emphasize that this meta-analysis procedure is modular with regard to choice of predictor, so we can substitute more sophisticated predictors as they become available or as is appropriate for particular phenotypes.

In order to achieve maximum power gain without sharing genotype data, a meta-analysis consortium would need to employ a iterative computational scheme^1^, which would require multiple exchanges of summary data between the cohorts and the meta-analysis group. Coordinating the submission of summary statistics to the central meta-analysis group is typically a bottleneck, so iterative approaches that require multiple rounds of data exchange will take much longer than a standard meta-analysis. Therefore, in this section we describe a simple one-step approach that provides an attractive compromise between speed and power.

The idea behind the fast, one-step design is that the meta-analysis consortium could construct a predictor using the summary statistics from the *last* meta-analysis. For example, the 2014 height study [2] could have used a predictor trained on the summary statistics from the 2010 height study [22], and the next height GWAS could use a predictor constructed from the 2014 height summary statistics. Similarly, the 2015 BMI study [3] could have used a predictor trained on the 2010 BMI study [23], and so forth.

The noncentrality parameter for this strategy is

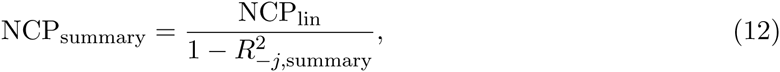

where 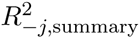 denotes the prediction *R*^2^ achieved by the predictor trained on summary statistics (leaving out the region around *j*).

There is no firm relationship between 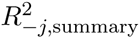 and 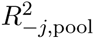. Our intuition is that when comparing equally sophisticated prediction algorithms, 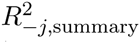 will typically be slightly lower than 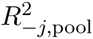, (which implies NCP_summary_ *<* NCP_pool_) because some information is lost when working with summary data instead and reference LD matrices instead of individual-level data and sample LD matrices [19].

Nevertheless, in typical scenarios where each *N*_*i*_ is one to two orders of magnitude smaller than *N*_meta_, we expect that the difference between 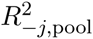 and *R*^2^ 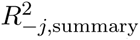 will be much less than the difference between 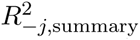 and 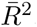 both 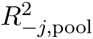 and 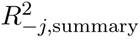 reflect the performance of predictors trained on a training set much larger than the training sets available to the predictors whose performance is measured by 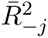.

Thus, the relationship between the noncentrality parameters is

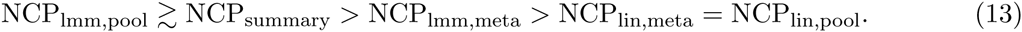

## Mixed Models for Sequencing Studies

The key idea behind the procedure in the previous section can be summarized in the following way: the predictions used in mixed model association testing do not need to be trained on the same sample used for testing. We should use the best genetic predictor (or combination of predictors) available, no matter how it is trained. This idea is also applicable to sequencing studies. At the time of writing, sequencing is roughly an order of magnitude more expensive than array genotyping; consequently, sequenced datasets tend to be roughly an order of magnitude smaller than genotyped datasets for the same trait. If we simply applied a mixed model (*e.g.,* BOLT-LMM) to the smaller sequenced datasets, the prediction *R*^2^ achieved by the mixed model predictor would be poor compared to what we could achieve by training a predictor on the larger genotyped datasets. Thus, we can increase power by using a predictor trained on the largest available genotyped dataset when running mixed-model association testing in the sequence data. Precisely,

1. Train a predictor on the SNP-array GWAS data. If individual-level genotype data are available, we can use the 

~~~
--predBetasFile
~~~

 flag in **bolt** to export prediction weights. If not, we can use LDpred to convert summary statistics into valid prediction weights.
2. Evaluate this predictor on the sequenced individuals (*e.g.,* using the 

~~~
--score
~~~

 flag in 

~~~
plink
~~~

 [24, 25]).
3. Perform association testing on the sequenced dataset using bolt with the predictions from the previous step as covariates *e.g.,* using the --covarFile and --qCovarCol flags (again, leaving one chromosome out at a time in order to avoid proximal contamination).

The noncentrality parameter for this test is the same as in Equation 12; the gain in power vs standard in-sample mixed model association depends on the increase in prediction *R*^2^ gained by training a predictor on the SNP-array GWAS data. Note that the *p*-values reported by bolt are only asymptotically valid, so the calibration may be poor for variants with very low minor allele count.

## Acknowledgements

Thanks B Neale, A Price, P Loh and B Vilhjalmsson and L Souchong for helpful comments.

## Footnotes

1 There are algorithms from machine learning that allow one to train a predictor by computing on only a subset of the data at any given time (*e.g.,* stochastic gradient descent and its more sophisticated parallel cousins). In machine learning, the barrier to holding all data in memory is typically data size, rather than privacy concerns, but the algorithms could nevertheless be adapted to GWAS consortia.

